# Three-dimensional Shear Wave Elastography Using Acoustic Radiation Force and A 2-D Row-Column Addressing (RCA) Array

**DOI:** 10.1101/2023.05.18.541365

**Authors:** Zhijie Dong, U-Wai Lok, Matthew R. Lowerison, Chengwu Huang, Shigao Chen, Pengfei Song

## Abstract

Acoustic radiation force (ARF)-based shear wave elastography (SWE) is a clinically available ultrasound imaging mode that noninvasively and quantitatively measures tissue stiffness. Current implementations of ARF-SWE are largely limited to 2-D imaging, which does not provide robust estimation of heterogeneous tissue mechanical properties. Existing 3-D ARF-SWE solutions that are clinically available are based on wobbler probes, which cannot provide true 3-D shear wave motion detection. Although 3-D ARF-SWE based on 2-D matrix arrays have been previously demonstrated, they do not provide a practical solution because of the need for a high channel-count ultrasound system (e.g., 1024-channel) to provide adequate volume rates and the delicate circuitries (e.g., multiplexers) that are vulnerable to the long-duration “push” pulses. To address these issues, here we propose a new 3-D ARF-SWE method based on the 2-D row-column addressing (RCA) array which has a much lower element count (e.g., 256), provides an ultrafast imaging volume rate (e.g., 2000 Hz), and can withstand the push pulses. In this study, we combined the comb-push shear elastography (CUSE) technique with 2-D RCA for enhanced SWE imaging field-of-view. *In vitro* phantom studies demonstrated that the proposed method had robust 3-D SWE performance in both homogenous and inclusion phantoms. An *in vivo* study on a breast cancer patient showed that the proposed method could reconstruct 3-D elasticity maps of the breast lesion, which was validated using a commercial ultrasound scanner. These results demonstrate strong potential for the proposed method to provide a viable and practical solution for clinical 3-D ARF-SWE.

## Introduction

Ultrasound shear wave elastography (SWE) is a clinically available imaging method that directly and quantitatively measures tissue stiffness [1-3]. SWE utilizes ultrafast ultrasound to track the fast-propagating shear wave motion. The shear wave speed is associated with tissue viscoelasticity, which is an essential biomarker for clinical diagnoses such as thyroid nodules [4, 5], liver fibrosis [6, 7], and breast cancer [8]. Shear wave motion can be induced by either external vibration [9, 10] or acoustic radiation force (ARF) [11, 12].

Similar to the mainstream ultrasound imaging techniques, current SWE methods are primarily confined to 2-D imaging, which does not provide a comprehensive analysis of the shear wave signal which propagates in all three dimensions. As a result, the reliability of the quantitative elasticity measurements may be compromised because of missing the shear wave signals that are not captured in the 2-D field-of-view (e.g., out-of-plane shear wave signals). Although for homogeneous tissues, the lack of 3-D shear wave tracking may not result in unreliable SWE performance, for anisotropic tissue or a complex shear wave field (e.g., external vibration), 3-D shear wave detection becomes indispensable [13-17].

Conventional 3-D SWE techniques are mainly based on mechanically tilting, rotating, or translating of 1-D array transducers (e.g., wobbler) to stack 2-D SWE slices into 3-D volumes [18, 19]. However, these SWE techniques may not be considered true 3-D because they do not detect shear wave propagation in all three dimensions. As such, these techniques do not provide unbiased 3-D shear wave speed estimation that gives rise to 3-D quantifications of shear elasticity. In addition, wobbler-based 3-D SWE takes several seconds to scan the 3-D volume, making it susceptible to handheld probe movement and tissue motion.

Over the past decade, 2-D matrix arrays-based 3-D SWE methods enabled high volume rate acquisition as well as 3-D motion detection [13, 20-22]. Because the whole volume can be rapidly scanned with electronic steering, 2-D matrix arrays are capable of detecting 3-D shear wave motion for 3-D shear elasticity estimation. However, one limitation of 2-D matrix arrays is that a high channel-count ultrasound system (e.g., 1024-channel for a 32×32 element 2-D array) is required to provide an adequate volume rate for 3-D shear wave tracking. This requirement imposes a practical barrier to the widespread adoption of the technology because high channel-count ultrasound systems are very expensive and not commonly available. A common solution for this issue is to use multiplexing [23, 24] and micro-beamforming [25, 26], which effectively reduces the requirement of a high channel-count ultrasound system. However, multiplexing reduces the imaging volume rate, which makes it challenging to track 3-D shear wave motion [21]. In addition, the complex circuitry associated with multiplexing and micro-beamformers is vulnerable to the long-duration push pulses used in ARF-SWE.

Recently, 2-D row-column addressing (RCA) arrays have emerged as an enticing solution for addressing the issues of low volume rate and high channel count with 2-D matrix arrays. 2-D RCA arrays do not use fully populated 2-D matrices for element distribution – instead, it uses bar-shaped elements arranged orthogonally as row elements and column elements [27], as shown in Fig. 1(a). As a result, the number of elements is reduced from *N*×*N* to *N*+*N*(e.g., 16,384 to 256 elements), making them compatible with mainstream ultrasound systems. This design makes RCA arrays advantageous for 3-D SWE for several reasons: First, the significantly reduced element count eliminates the need for multiplexing and micro-beamforming, which makes 2-D RCA arrays suitable for generating ARF-based shear waves with long-duration push pulses; Second, when combined with a system that is equipped with software beamformers and massive parallel receive capability (e.g., capable of plane wave imaging), 2-D RCA arrays provide an ultrafast 3-D imaging volume rate (e.g., several thousand Hertz) that is ideal for 3-D shear wave tracking. In a previous study [28], we validated RCA-based 3-D shear wave detection by using both mechanical vibrations and ARF (by using a separate probe). In another study, Bernal *et al* used RCA arrays to detect passive shear waves [29]. To date, a 3-D ARF-SWE approach based on the same 2-D RCA arrays (i.e., for both shear wave generation and detection) has not been reported, which is what we will present in this study.

**Fig. 1.**
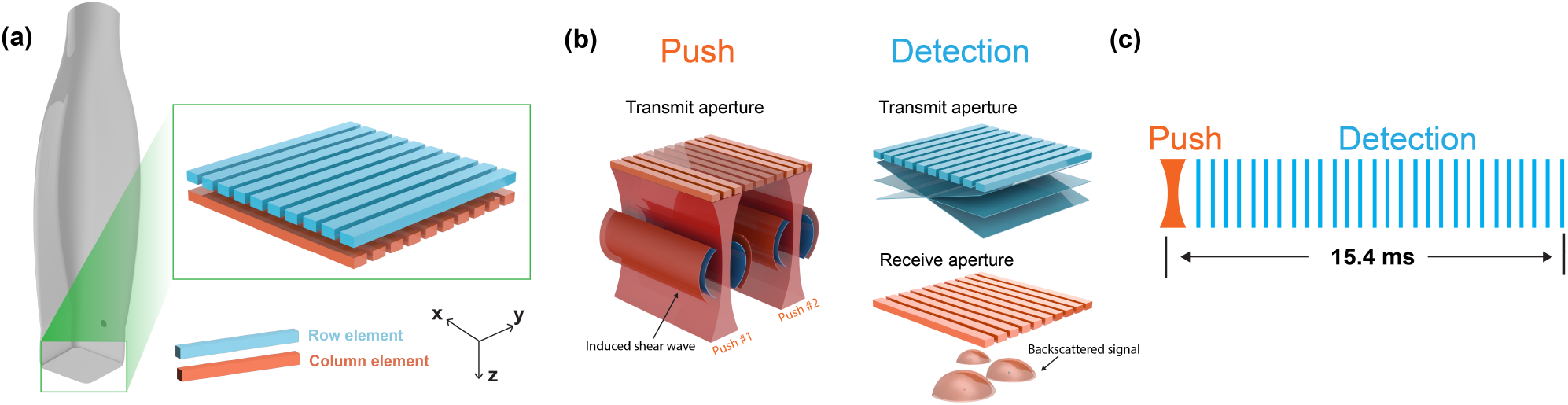
(a) Row-column addressing (RCA) array with orthogonally arranged row elements (distributed along the x-direction) and column elements (distributed along the y-direction). (b) Illustration of the push and detection schemes of the 3-D ARF-based SWE combined with comb-push shear elastography (CUSE). The column elements transmit two focused push beams to generate shear wave motion, followed by shear wave detection using the row elements for transmit focusing and column elements for receive focusing. (c) Timeline of the SWE sequence. The two push beams (400 *μS* each) were simultaneously (focused comb-push) or successively (marching comb-push) transmitted at first, followed by an ultrafast acquisition with 30 volumes at a 2000 Hz volume rate. The total duration of push-detection data acquisition was 15.4 ms for focused comb-push.

The rest of this paper is structured as follows. We first describe the workflow and sequences of the proposed method, followed by phantom studies and an *in vivo* case study on a breast cancer patient. The corresponding results will be presented in Section III. We finalize this article with discussions and conclusions.

## Materials and Methods

### Implementation and Optimization of 3-D ARF-SWE based on the 2-D RCA Array

A custom-built RCA array (Daxsonics Ultrasound Inc., Halifax, Canada; central frequency at 7 MHz with 65% bandwidth) and a Verasonics Vantage 256 system (Verasonics Inc., Kirkland, WA) were used in this study for ARF-based shear wave generation and 3-D shear wave detection. The RCA array includes 128 row elements stacked on top of 128 column elements (Fig. 1(a)). Thanks to the simple design, the 2-D RCA array can sustain the long push pulses for ARF generation (e.g., hundreds of microseconds). The configurations of the RCA array as well as the 3-D SWE imaging sequences are reported in Table I.

**Table I.**
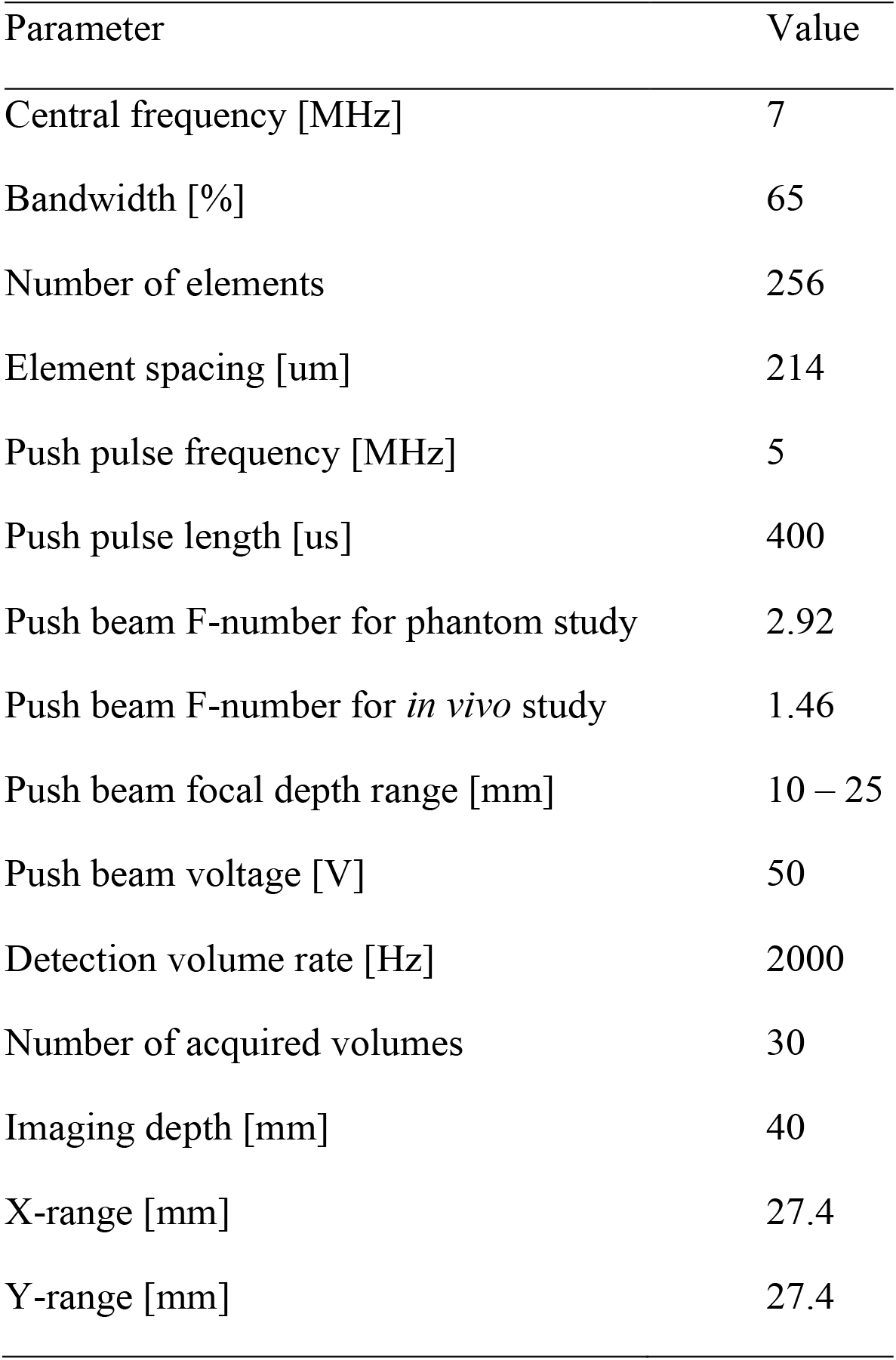
3-D ARF-SWE Imaging Configurations

In this study, we combined 2-D RCA with the comb-push shear elastography (CUSE) technique to achieve a large 3-D imaging field-of-view (FOV) without the need for multiple push-detection data acquisitions [30]. As shown in Fig. 1(b), two focused push beams positioned at the edges of the transducer were simultaneously (focused comb-push) or successively (marching comb-push) [31] transmitted for shear wave generation using column elements (distributed along the y-direction). Because of the bar-shaped element, 2-D RCA arrays can only perform transmit focusing in one dimension, which results in plate-shaped shear waves that are focused in one dimension and planar in the orthogonal dimension (Fig. 1(b)). Various push beam configurations were tested, and a final configuration of a push pulse with 400 us duration, and 5 MHz center frequency was used throughout the rest of the study, as detailed in Table I.

For 3-D shear wave detection, a compounding plane wave imaging sequence with 7 compounding angles was used immediately after the push beam transmission, as shown in Fig. 1(b) and (c). The angular step size of 0.5 degrees was used to mitigate grating lobes [32]. For detection, all row elements (distributed along the x-direction) were used to transmit, and all column elements were used to receive. As a result, the transmit focusing was along the x-direction, and the dynamic receive focusing was along the y-direction, which was aligned with the main propagation direction (y-direction) of the induced shear waves. A total of 30 volumes were acquired at a 2000 Hz volume rate. The final 3-D SWE sequence (including a single push and detection cycle) for the entire 3-D volume data acquisition only lasted 15.4 ms, which is significantly faster than existing 3-D SWE methods (e.g., a few seconds for wobbler-based 3-D SWE [18], and around 128 ms for matrix array-based 3-D SWE with 5 push cycles and 1488 Hz volume rate [20]). Throughout the studies, a B-mode imaging sequence (50 Hz volume rate, 31 compounding angles) with real-time bi-planar visualization of 3-D volumes was used to provide scanning guidance and localize the targeted tissue, followed by the 3-D ARF-SWE acquisition.

For 3-D shear wave speed (SWS) map reconstruction, the following processes were implemented, as outlined in Fig. 2. First, acquired in-phase quadratic-phase (IQ) volumes were beamformed using delay-and-sum (DAS) beamforming (Fig. 2(a)), followed by raw shear wave motion estimation (Fig. 2(b)) using 1-D autocorrelation [33, 34]. Subsequently, bandpass filtering and directional filtering were applied to denoise and separate the shear wave signal into two directions (positive y-direction and negative y-direction), as shown in Fig. 2(c). Finally, the 3-D cross-correlation [20, 35] method was used to reconstruct two 3-D SWS maps for each directional filter, which were then combined into a final 3-D SWS map using the Tukey window [36], as shown in Fig. 2(d).

**Fig. 2.**
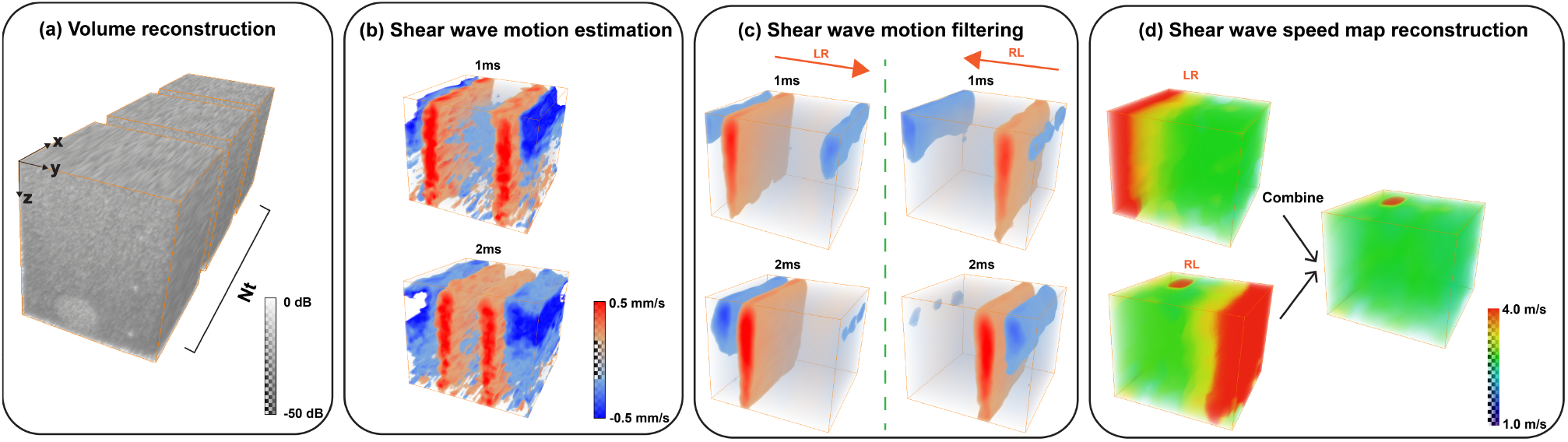
The proposed 3-D SWE data processing steps. (a) IQ volumes acquired using the RCA array at an ultrafast volume rate (e.g., 2000 Hz) were beamformed. (b) Raw shear wave motions were calculated from acquired IQ volumes. (c) Filtered shear wave motions from left to right (LR) and from right to left (RL) were extracted using directional filtering and bandpass filtering. (d) 3-D SWS maps from separated shear wave fields were reconstructed and combined to reconstruct the final 3-D SWS map.

### Acoustic Field Scan and Acoustic Output Measurements

The acoustic field and output for the push beams used in this study were assessed using both simulation and water tank measurements with hydrophones. The simulation was conducted using the Verasonics simulator. For the hydrophone measurement, a capsule hydrophone (HGL-0200, Onda Corp, Sunnyvale, CA), submerged in a water tank filled with deionized and degassed water using a water conditioner (AQUAS-10, Onda Corp, Sunnyvale, CA), was used to measure the 3-D acoustic pressure field of the push beams. The hydrophone was mounted on linear and rotary positioners (AIMS III, Onda Corp, Sunnyvale, CA) to scan the 3-D acoustic field, with 1-, 1-, and 2-mm step size in the x (lateral), y (elevational), and z (axial) dimensions, respectively. The Vantage system was synchronized with the capsule hydrophone and the scanning stage, and the water temperature was measured to calibrate the speed of sound. MI (derated at a rate of 0.3 dB/cm/MHz) and *I*_*SPTA*,0.3_ were measured for the push beams with different focal depths (10-25 mm) and F-numbers (1.46 and 2.92).

### Elasticity Phantom Studies

The performance of the proposed 3-D ARF-SWE method was first evaluated in a multipurpose multi-tissue ultrasound phantom (Model 040GSE, CIRS Inc. Norfolk, VA, USA) and an elasticity phantom (Model 049, CIRS Inc. Norfolk, VA, USA). For the multi-purpose phantom, a homogenous region (shear wave speed of 2.60 ± 0.15 m/s [36]) was imaged to evaluate the accuracy of the shear wave speed measurement. For the elasticity phantom, four spherical lesion objects with different elasticity (SWS from 1.38 m/s to 4.65 m/s) were imaged to test the 3-D SWE performance in heterogeneous tissue. The targeted lesions were located at 10-20 mm depth with an approximate diameter of 10 mm, and a focal depth of 20 mm was used for the push beam with a F-number of 2.92. Both phantoms have an ultrasound attenuation of 0.5 dB/cm/MHz [37] and a speed of sound of 1540 m/s.

### In Vivo Case Study

To further evaluate the *in vivo* performance of the proposed 3-D ARF-SWE method, a case study on a breast cancer patient was performed. The same subject was also imaged by a conventional clinical ultrasound system (General Electric Healthcare, Wauwatosa, WI, USA) with 2-D SWE [30], which was used as a reference for the newly developed 3-D ARF-SWE method. The *in vivo* protocols were approved by the Mayo Clinic Institutional review board (IRB), and written informed consent was obtained before scanning. To improve the signal-to-noise ratio (SNR) of the shear wave signals, the F-number of the push beams was reduced to 1.46, and a marching comb-push with successive transmission of focused push beams was used. The total data acquisition time for the 3-D SWE sequence was approximately 16 ms.

## Results

### Acoustic Field Scan and Acoustic Output Measurements

Fig. 3 shows the simulated and measured 3-D acoustic fields of the comb-push beams, as well as two representative slices along the y-z and x-y dimensions. The measured fields were closely matched with the simulation results. Due to the element sensitivity difference resulting from probe manufacturing, the acoustic fields of the left push beam and right push beam were not identical.

**Fig. 3.**
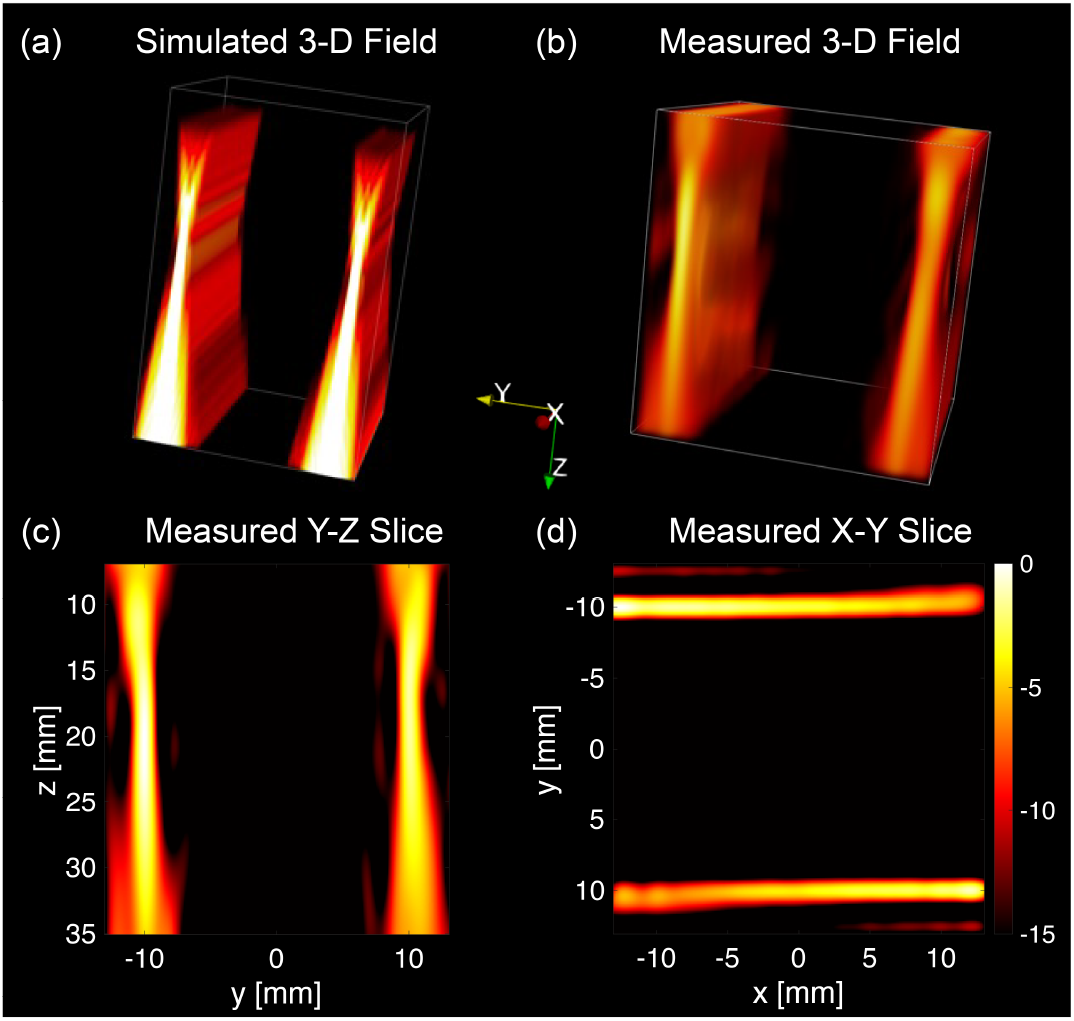
Simulated and measured 3-D acoustic fields of the comb-push beams generated by the RCA array. (a) Simulated 3-D acoustic field of the comb-push beams. (b) Hydrophone-measured 3-D acoustic field. (c) Measured acoustic beam in the y-z plane at x = 0 mm. (d) Measured acoustic beam in the x-y plane at z = 20 mm. A dynamic range of 15 dB was used.

The MI values measured from 5 mm to 30 mm for push beams with two different F-numbers (i.e., 2.92 and 1.46) and different focal depths were plotted in Fig. 4. A maximum MI value of 1.23 was measured when the push beam was focused at 15 mm depth with an F-number of 1.46, and the maximum *I*_SPTA,0.3_ measured was 132.09 mW/cm^2^ with a 1 Hz PRF, which are both well below the Food and Drug Administration (FDA) regulatory limit [38].

**Fig. 4.**
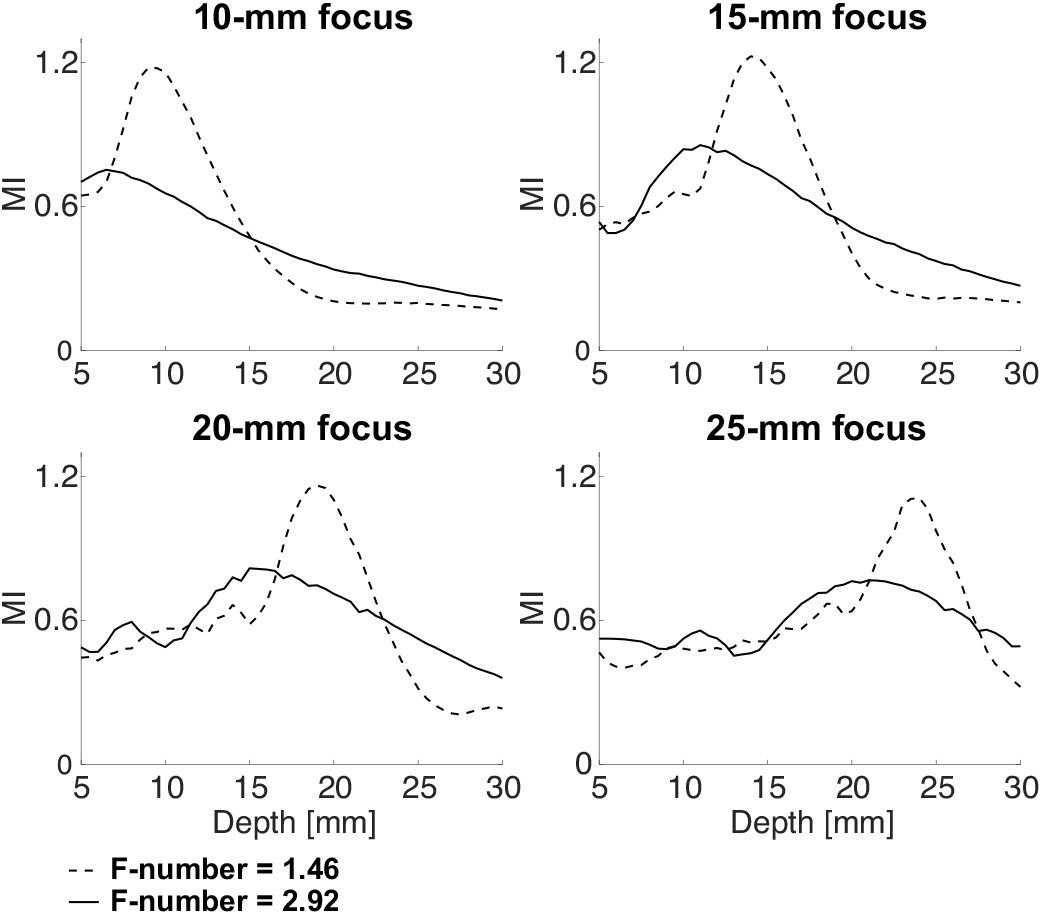
MI measurements of the push beams with different focal depths (10-25 mm) and different F-numbers (1.46 and 2.92).

### Elasticity Phantom Studies

Fig. 5(a) presents the representative 3-D shear wave motions induced by the comb-push beams using the RCA array in the homogenous phantom. Shear waves were successfully induced by ARF and propagated across the full FOV in two opposite directions (positive y-direction and negative y-direction). Fig. 5(b) shows the reconstructed 3-D SWS map of the homogenous phantom, as well as the histogram. The measured SWS was 2.63 ± 0.13 m/s, which was well matched with the reference value (2.60 ± 0.15 m/s) from the literature [36]. The results from the *in vitro* homogenous phantom study demonstrated that the proposed method could successfully perform 3-D shear wave detection with ultrafast volume rate (e.g., 2000 Hz) and reconstruct the full 3-D SWS map with high accuracy.

**Fig. 5.**
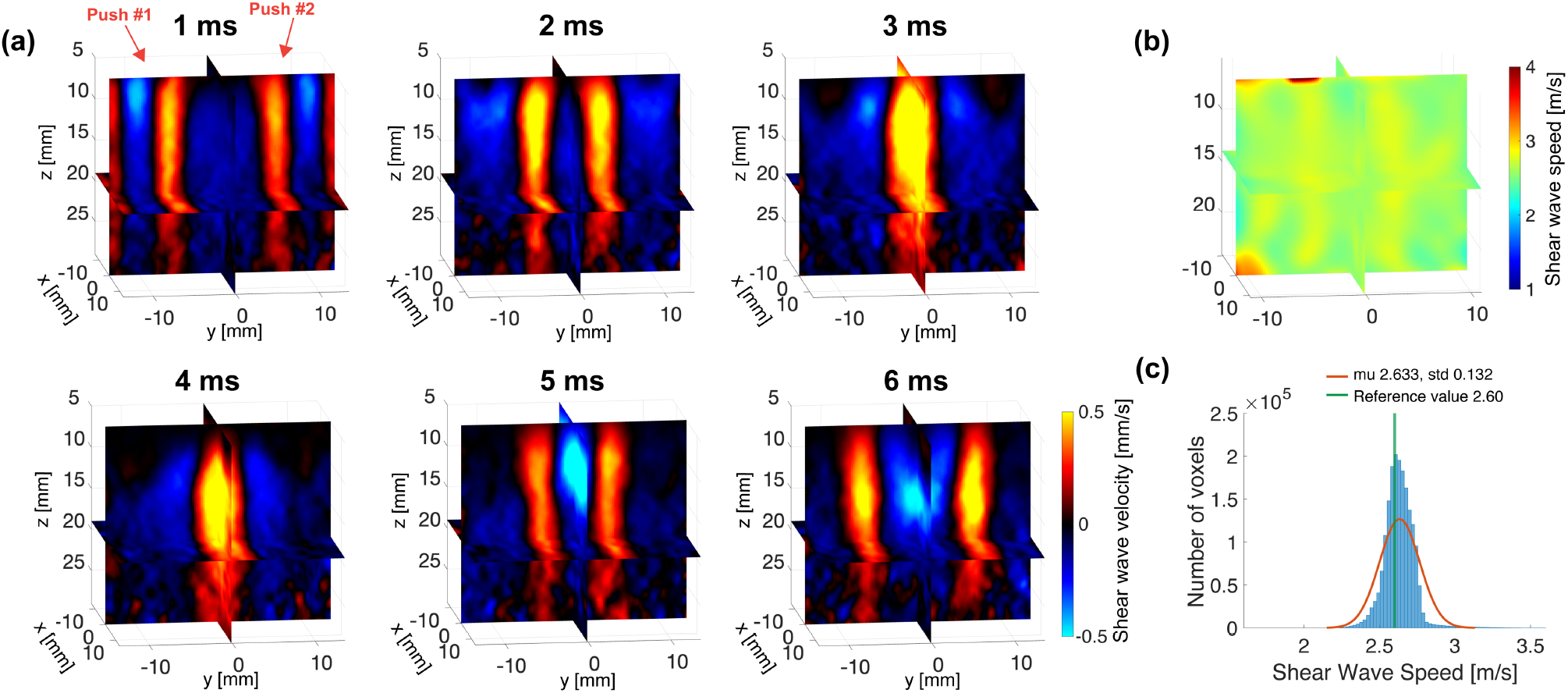
3-D shear wave motions and reconstructed 3-D SWS map of the homogenous phantom using the proposed 3-D SWE method. (a) Tri-plane view of the 3-D shear wave motions induced by comb-push beams at different representative time points, and the detection volume rate was 2000 Hz. (b) Tri-plane view of the reconstructed 3-D SWS map. (c) SWS histogram of the full volume.

Fig. 6 demonstrates representative shear wave motions in the region containing a stiff spherical lesion (Type IV, 64.9 kPa) with higher SWS (4.65 m/s) compared to the background (2.45 m/s). As indicated by the arrows in Fig. 6, the shear waves accelerated when propagating through the lesion. The reconstructed 3-D SWS map is shown in Fig. 7, from which one can see the reconstructed SWS image was well aligned with the spherical lesion in the B-mode image [see Fig. 7(a) and (b)], and the lesion was clearly reconstructed in 3-D with a spherical shape (Fig. 7(c)). The measured volume of the segmented spherical lesion by thresholding (2.95 m/s) was 0.59 mL, which was matched with the nominal volume of the lesion (0.60 mL).

**Fig. 6.**
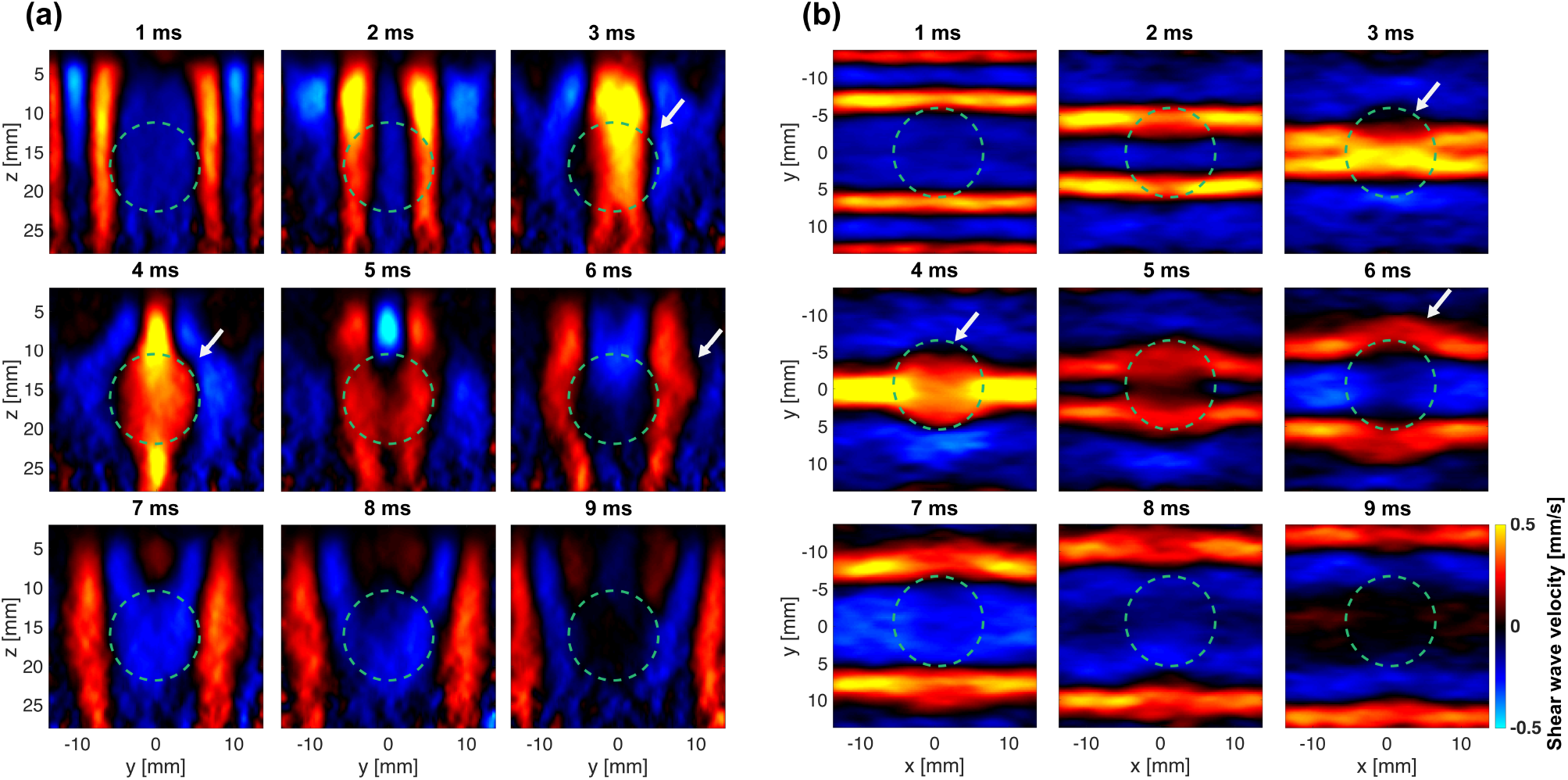
Shear wave motions at representative time points along the y-z plane (a) and x-y plane (b) in the elasticity phantom including a stiff spherical lesion object. The contour of the spherical lesion is indicated using the green dashed circle.

**Fig. 7.**
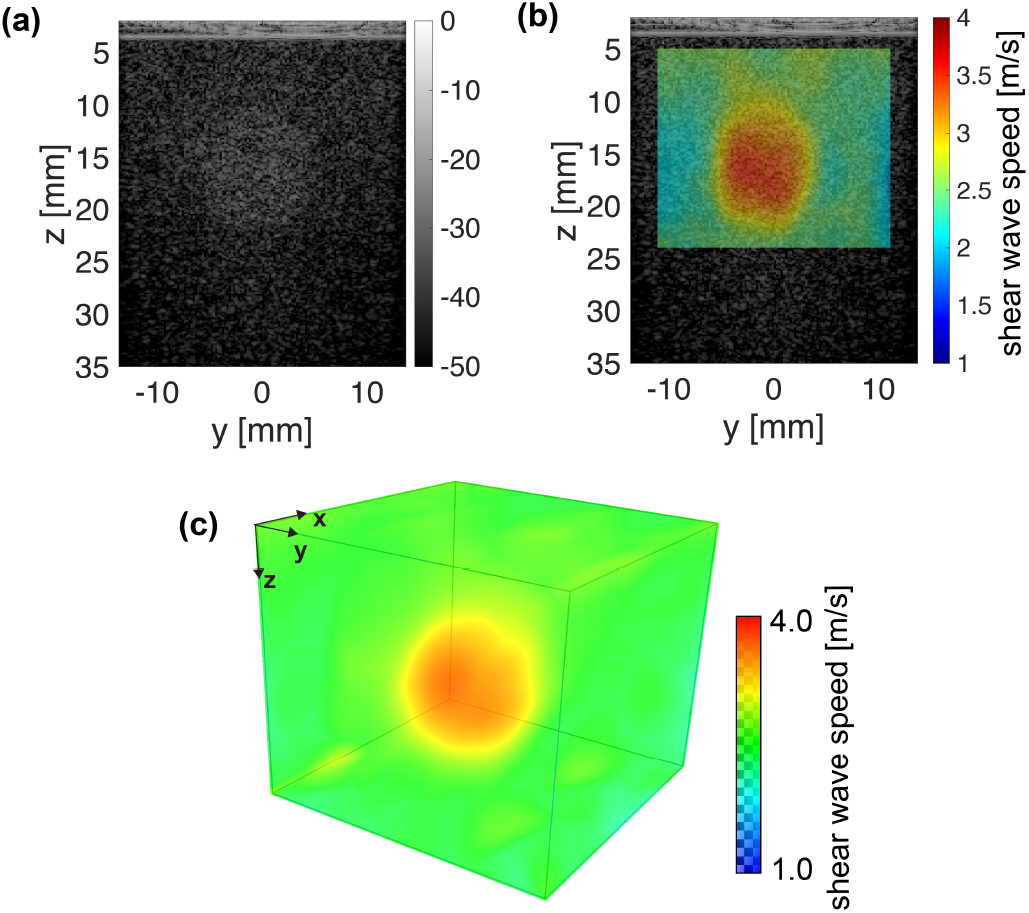
Reconstructed 3-D SWS map of the elasticity phantom, including a stiff spherical lesion object. (a) y-z slice of B-mode image by the RCA array, (b) y-z slice of the SWS map overlapped on the B-mode image, and (c) volumetric view.

Fig. 8 shows the 3-D SWS results of four different types of lesions. The spherical lesions with different shear wave speed values can be clearly visualized (Fig. 8(a)). As shown in Fig. 8(b), for lesion type I, the reference SWS was 1.38 m/s, and the measured SWS was 1.86 ± 0.05 m/s; for lesion type II, the reference SWS was 1.83 m/s, and the measured SWS was 2.00 ± 0.03 m/s; for lesion type III, the reference SWS was 3.45 m/s, and the measured SWS value was 2.72 ± 0.03 m/s; for lesion type IV, the reference SWS was 4.65 m/s, and the measured SWS value was 3.37 ± 0.09 m/s; for background, the reference SWS was 2.45 m/s, and the measured SWS value was 2.36 m/s ± 0.06 m/s. The quantitative measurements show that the proposed method can differentiate different lesions based on measured shear wave speeds. However, the calculated SWS values were overestimated for soft lesions and underestimated for stiff lesions, the biased measurements may be due to the partial volume effect and spatial resolution limit of the proposed method, which resulted from the limited number of compounding angles for high volume rate and RCA layout (i.e., only one-way focusing along the x- and y-direction).

**Fig. 8.**
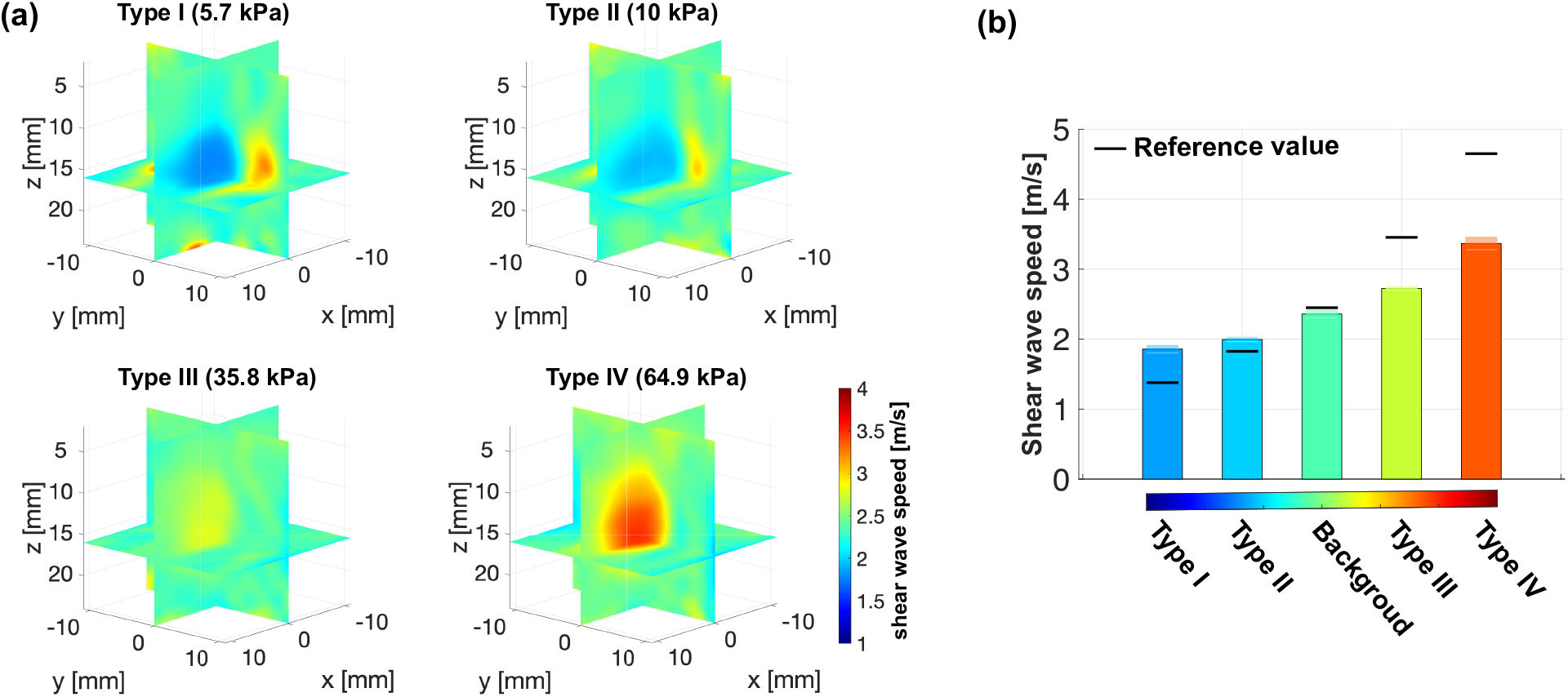
Reconstructed SWS maps and calculated SWS values of four different types of spherical lesions. (a) Tri-plane view of reconstructed SWS maps. Corresponding Young’s modulus are labeled. (b) Calculated SWS values of four types of lesion objects and background of the elasticity phantom. The corresponding standard deviation and reference values (black line) are labeled.

### In Vivo Case Study

Fig. 9 shows the *in vivo* B-mode images and SWS images in the breast of a volunteer using clinical 2-D ultrasound systems and the proposed 3-D ARF-SWE method. The B-mode image from the conventional clinical scanner shows a breast mass, and the corresponding 2-D SWS image indicates a mass with high stiffness [see Fig. 9(a) and (b)]. The RCA array shows similar results with a stiff lesion. The reference SWS measurement using the 2-D SWE from the conventional clinical system was 6.60 m/s, and the measured SWS using the proposed method was 8.43 m/s. The higher SWS value from RCA may be a result of probe compression during scanning, which was necessary because of the suboptimal B-mode imaging quality of the RCA.

**Fig. 9.**
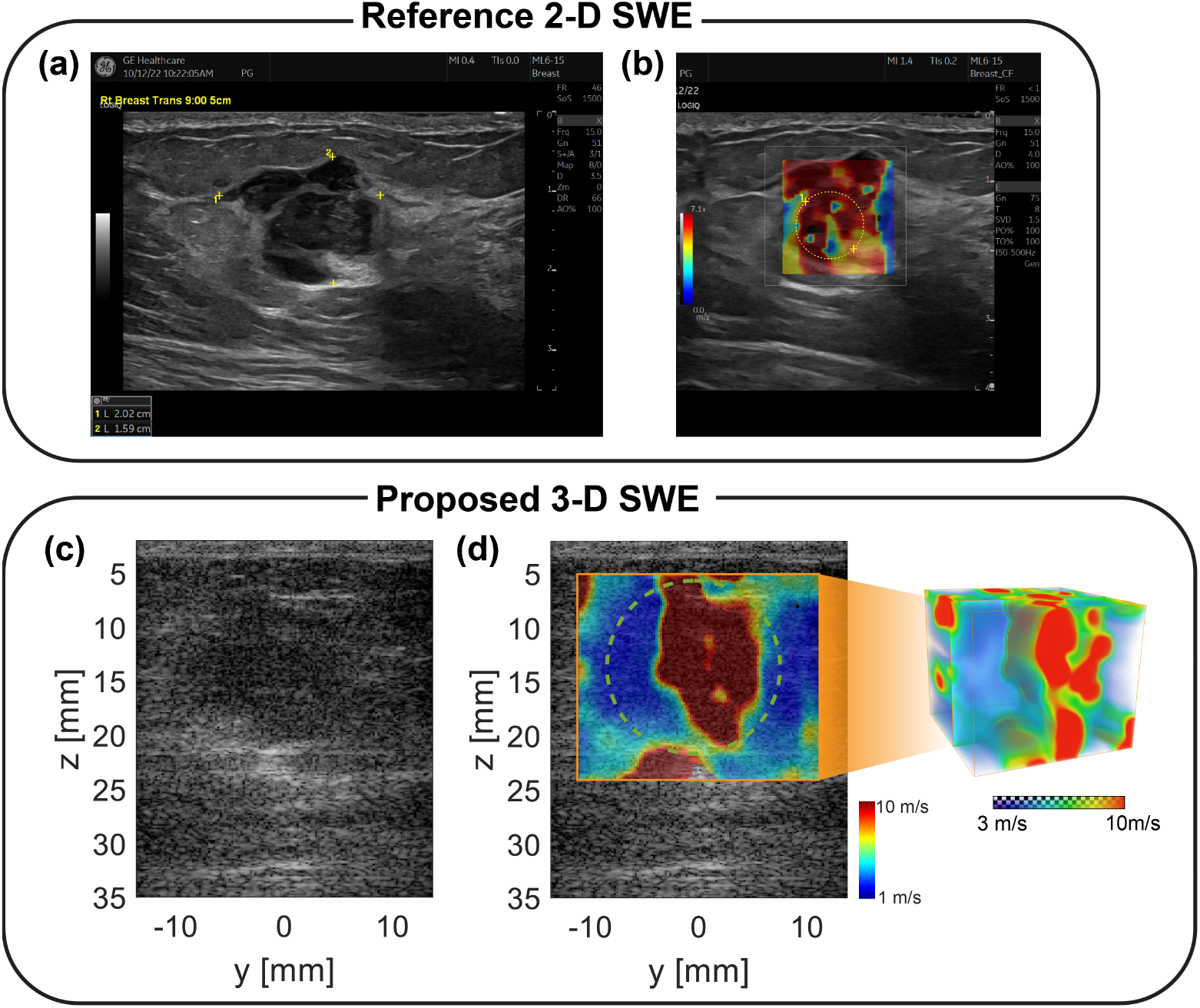
B-mode and SWE imaging of a breast lesion with the reference 2-D SWE method and the proposed 3-D ARF-SWE method. (a) Conventional B-mode image of a breast mass using the clinical scanner. (b) 2-D SWE using the clinical scanner. (c) B-mode image along the y-z slice using the RCA array. (d) Reconstructed 3-D SWS map and the y-z slice using the proposed 3-D SWE method. The SWS measurement ROIs are labeled in (b) and (d).

## Discussions

Previously we developed an RCA-based 3-D shear wave motion detection technique that utilized external vibration to perform 3-D SWE [28]. RCA arrays provide an ultrafast volume rate (e.g., 2000 Hz) with a short acquisition time (e.g., tens of milliseconds), which is ideal for practical implementations of the 3-D SWE technology. In this study, we further developed the 3-D SWE technique to perform ARF-based SWE using the same RCA probe. We incorporated the comb-push shear elastography (CUSE) technique in our study to achieve fast and full FOV 3-D SWS reconstruction. In this study, we demonstrated that the proposed method could reconstruct a 3-D SWE map using a single push-detect data acquisition (15.4 ms) at an ultrafast volume rate (e.g., 2000 Hz). The acoustic field scanning illustrated that the RCA array could generate robust push beams with adequate ARF. The RCA was able to withstand the long-duration push pulses without causing any damage to the probe. The quantitative and qualitative results from elasticity phantom studies and an *in vivo* study demonstrated that the 3-D shear wave motions and 3-D elasticity map could be successfully detected and reconstructed by the proposed method, respectively. Furthermore, the *in vivo* breast imaging results showcase the human imaging capacity of our method, indicating a clear path toward clinical implementation.

Compared to the external vibration-based 3-D SWE, the ARF-based 3-D SWE method is more efficient and convenient because there is no need for additional equipment (e.g., vibrators and related electronic equipment), and push and detection can be performed using only the RCA array. Furthermore, compared to the low frequencies used by the external vibration (e.g., harmonic within the range of 50 – 200 Hz), the shear waves induced by the ARF have a broader spectrum (e.g., 75-500 Hz [39]), corresponding to smaller shear wave wavelength and better spatial resolution for the reconstructed SWS map.

One significant advantage of the proposed method compared to the clinical 3-D SWE technologies is that the proposed method could perform true 3-D shear wave detection with sufficient acoustic radiation force at ultrafast volume rate (e.g., thousands of Hertz). The conventional 3-D SWE technique is based on the mechanical translation of the 1-D array (e.g., wobblers), which limits the shear wave detection in 2-D planes, and the inter-plane shear wave motions cannot be retrieved. For 3-D SWE methods based on 2-D matrix arrays using ARF, the FOV is largely limited (e.g., 19.2 × 19.2 × 20 *mm*^3^ for a 32 × 32 array with 1024 channels [20]), and increasing the number of channels (e.g., several thousand) for enlarged FOV is technically challenging, which also restricts their clinical feasibility. Furthermore, as the multiplexer is necessary to reduce the number of physical channels in 2-D matrix arrays, the volume rate is significantly reduced (e.g., hundreds of Hertz), and the delicate circuitry related to the multiplexers is susceptible to damage caused by the long-duration push pulses. Thanks to the unique design of the RCA array, the proposed method with shear wave generation and 3-D detection capabilities is compatible with clinical ultrasound systems and has a clear pathway for future clinical translation.

Another benefit of the proposed method is that the whole 3-D elasticity map reconstruction can be done using a single push event with the comb-push beams. One issue related to the ARF-based SWE is that the shear wave speed cannot be retrieved in the focused beam region. By simultaneously transmitting two push beams from both sides of the RCA array, the focal region of the push beam is covered by the shear waves induced by the other push beam from the opposite side, as shown in Fig. 2. The proposed method used comb-push beams (0.4 ms) and continuous acquisition of 30 volumes (15 ms) at 2000 Hz, leading to a total acquisition time of 15.4 ms, which was fast enough to avoid tissue motions and handheld probe movement. Moreover, the patient did not need to hold their breath for several seconds during the data acquisition, which was a common practice for conventional 3-D SWE techniques. Note that number of acquired volumes needs to be increased for low shear wave speed estimation (e.g., 1 m/s), otherwise, the reconstruction FOV will be limited.

One limitation of the proposed method is related to the spatial resolution of RCA arrays, which is suboptimal because of the one-way focusing (e.g., either transmit or receive focusing along x-or y-direction, no two-way focusing in both dimensions). As an example, when transmitting using row elements (distributed along the x-direction) and receiving using column elements (distributed along the y-direction), as indicated in Fig. 1, the transmit focusing is along the x-direction, and the receive focusing is along the y-direction, which is different from two-way focusing of the 2-D matrix array. To achieve ultrafast volume rate (i.e., thousands of Hertz) as required for ARF-induced shear wave detection without aliasing, the number of compounding angles is limited (e.g., 7 angles used in this study), resulting in suboptimal spatial resolution along x-direction as shown in Fig. 1(b), where only transmit focusing was available, and therefore the spatial resolution is strictly dictated by the number and range of compounding angles [40]. Therefore, the spatial resolution along the transmit focusing of the reconstructed SWS map was worse than the one in the other direction, as shown in Fig. 8. This limitation can be alleviated by applying more compounding angles or using other transmit schemes such as synthetic aperture [41]; however, the volume rate will be reduced, and multiple push pulse transmissions might be needed to synthesize a high volume rate acquisition.

The imaging quality of 3-D elasticity maps depends on two major factors: shear wave signal quality (e.g., shear wave SNR and spectrum, which is largely dictated by the power of the push beam) and shear wave detection imaging quality (e.g., ultrasound imaging SNR and spatial resolution). As discussed, the detection imaging quality is related to the intrinsic limitations associated with the RCA probe design. The phantom studies showed that the SWS measurements of the lesion objects were biased when compared to the reference SWS values, which could be resulted from the partial volume effect and suboptimal spatial resolution related to the RCA array. Ongoing studies are being conducted to further optimize the sequence and processing to address the biasing issue.

Although the current RCA array provides sufficient acoustic radiation force for inducing shear waves, its MI is lower than conventional 1-D probes (e.g., MI of 1.5 or higher [28]). As a result, the generated shear waves are weaker than conventional 1-D ultrasound, posing a challenge for clinical studies. To achieve higher MI values for strong shear wave signals, F-number can be reduced with more activated elements at each transmission. Furthermore, multiple push events can be used to improve SNR and contrast, for example, comb-push beams could be applied multiple times with different push locations, and then compound to the final elastography image. Meanwhile, a better RCA probe design and manufacturing is necessary to improve the quality of the probes, and further work can be conducted to optimize the probe performance for better shear wave generation.

## Conclusions

In this study, we presented a novel 3-D SWE method using a 2-D row-column-addressing (RCA) array for both shear wave generation (i.e., using acoustic radiation force) and detection. The proposed 3-D SWE method provides 3-D shear wave speed maps with an ultrafast shear wave motion detection volume rate (e.g., 2000 Hz). We integrated the comb-push shear elastography (CUSE) technique into our approach, resulting in a fast 3-D SWE sequence with a total data acquisition time of approximately 16 milliseconds. *In vitro* and *in vivo* results demonstrated robust performance of the proposed 3-D SWE technique, which provides a practical and cost-effective 3-D SWE solution for clinical implementations of 3-D tissue elasticity imaging.

## Acknowledgment

This work was supported by the Department of Defense (DoD) through the Breast Cancer Research Program (BCRP) under Award Nos. W81XWH-21-1-0062 and W81XWH-21-1-0063. Opinions, interpretations, conclusions, and recommendations are those of the author and are not necessarily endorsed by the Department of Defense.

